# A universal formula for avian egg shape

**DOI:** 10.1101/2020.08.15.252148

**Authors:** Valeriy G. Narushin, Michael N. Romanov, Darren K. Griffin

## Abstract

The bird’s oomorphology has far escaped mathematical formulation universally applicable. All bird egg shapes can be laid in four basic geometric figures: sphere, ellipsoid, ovoid, and pyriform (conical/pear-shaped). The first three have a clear mathematical definition, each derived from expression of the previous, but a formula for the pyriform profile has yet to be inferred. To rectify this, we introduced an additional function into the ovoid formula. The subsequent mathematical model fits a completely novel geometric shape that can be characterized as the last stage in the evolution of the sphere—ellipsoid—Hügelschäffer’s ovoid transformation applicable to any avian egg shape geometry. Required measurements are the egg length, maximum breadth, and diameter at the terminus from the pointed end. This mathematical description is invariably a significant step in understanding not only the egg shape itself, but how and why it evolved, thus making widespread biological and technological applications theoretically possible.

## Introduction

Described as “the most perfect thing” (***Birkhead, 2016***), the avian egg is one of the most recognizable shapes in nature. Despite this, an expression of “oviform” or “egg-shaped” (a term used in common parlance) that is universally applicable to all birds has belied accurate description by mathematicians, engineers and biologists (***Narushin et al., 2020a***). Various attempts to derive such a standard geometric figure in this context that, like many other geometric figures, can be clearly described by a mathematical formula are nonetheless over 65 years old (***Preston, 1953***). Such a universal formula potentially has applications in disciplines such as evolution, genetics, ornithology, species adaptation, systematics, poultry breeding and farming, food quality, engineering, architecture and artwork where oomorphology (***Mänd et al., 1986***) is an important aspect of research and development.

According to ***Nishiyama*** (***2012***), all profiles of avian eggs can be described in four main shape categories (1) *circular, elliptical, oval* and *pyriform* (conical/pear-shaped). A circular profile indicates a spherical egg; elliptical an ellipsoid; oval an ovoid and so on.

Many researchers have identified to which shape group a particular egg can be assigned, and thus developed various indices to help make this definition more accurate. Historically, the first of these indices was the shape index (*SI*) ***Romanoff and Romanoff*** (***1949***), which is a ratio of maximum egg breadth (*B*) to its length (*L*). *SI* has been mainly employed in the poultry breeding industry to evaluate the shape of chicken eggs and sort them thereafter. Its disadvantage is that, according to this index, one can only judge whether or not an egg falls into the group of circular shape. With each subsequent study, there have been more and more other devised indices. That is, while the early studies (***Preston, 1968***) limited themselves to the usefulness of such egg characteristics as asymmetry, bicone and elongation, the later ones increased the number of indices to seven (***Mänd et al., 1986***), and even to ten (***Mytiai and Matsyura, 2017***). The purpose of the current study was to take this research to its ultimate conclusion to present a universal formula for calculating the shape of any avian egg based on revising and re-analysis of the main findings in this area.

In parallel to the process of developing various egg shape indices, a broader mathematical insight into comprehensive and optimal description of the natural diversity of oviform warrants further study. The definition of the groups of circular and elliptical egg shapes (***Figure 1A–B***) is relatively straightforward since there are clear mathematical formulae for the circle and ellipse. To describe mathematically oval and pyriform shapes (***Figure 1C–D***) however, new theoretical approaches are necessary.

**Figure 1.**
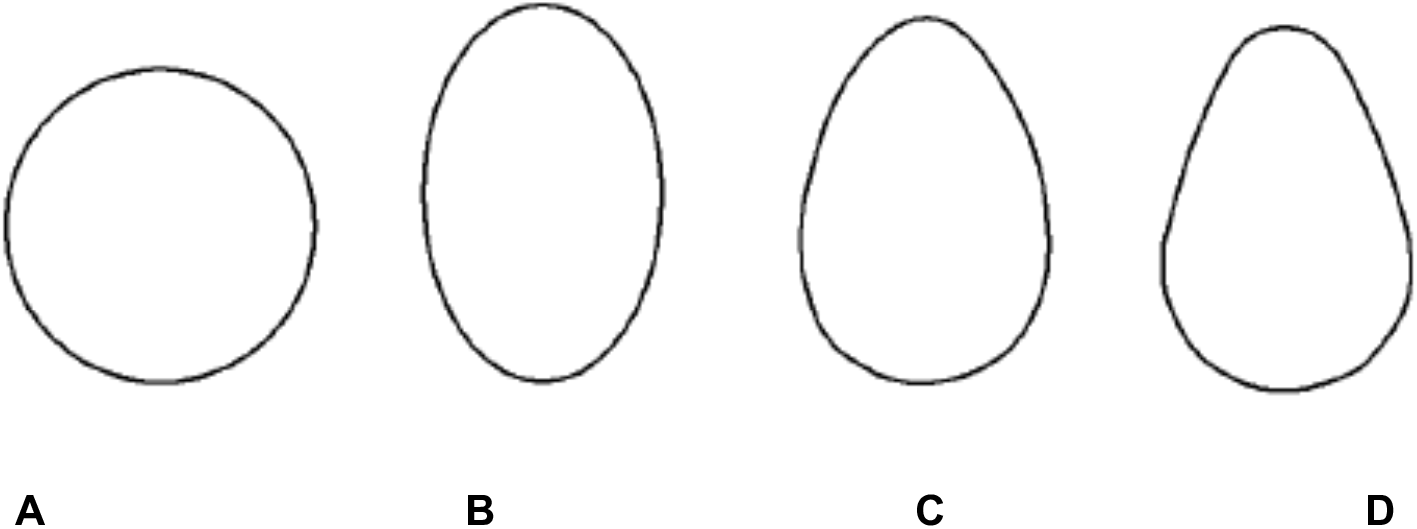
Basic egg shape outlines based on ***Nishiyama*** (***2012***): (**A**) circular, (**B**) elliptical, (**C**) oval, and (**D**) pyriform.

***Preston*** (***1953***) proposed the ellipse formula as a basis for all egg shape calculations. Multiplying the length of its vertical axis by a certain function *f*(*x*) (which he suggested to express as a polynomial) Preston showed that most of the eggs studied could be described by a cubic polynomial, although for some avian species, a square or even linear polynomial would suffice. This mathematical hypothesis turned out to be so effective that most of the further research in this area was aimed solely at a more accurate description of the function *f*(*x*). Most often, this function was determined by directly measuring the tested eggs, after which the data was subjected to a mathematical processing using the least squares method. As a result, a function could be deduced that, unfortunately, would be adequate only to those eggs that were involved in an experiment (***Baker, 2002; Troscianko, 2014; Pike, 2019***). Some authors (***Todd and Smart, 1984; Biggins et al., 2018***) applied the circle equation instead of ellipse as the basic formula, but the principle of empirical determination of the function *f*(*x*) remained unchanged. Several attempts were made to describe the function *f*(*x*) theoretically in the basic ellipse formula (***Carter, 1968; Smart, 1991***); however, for universal and practical applicability to all avian eggs (rather than just theoretical systems), it is necessary to increase the number of measurements and the obtained coefficients.

The main problem of finding the most convenient and accurate formula to define the function *f*(*x*) is the difficulty in constructing graphically the natural contours corresponding to the classical shape of a bird’s egg (***Köller, 2000; Landa, 2013; Cook, 2018***). Indeed, all the reported formulae have a common flaw; that is, although these models may help define egg-like shapes in works of architecture and art, they do not accurately portray “real life” eggs for practical and research purposes. This drawback can be explained by the fact that the maximum breadth of the resulting geometric figure is always greater than the breadth (*B*) of an actual egg, as the *B* value is measured as the egg breadth at the point corresponding to the egg half length. This drawback has been reviewed in more detail in our previous work (***Narushin et al., 2020b***). In order, therefore, for the mathematical estimation of the egg contours not to be limited by a particular sample used for computational purposes, but to apply to all avian egg shapes present in nature, further theoretical considerations are essential. One such tested and promising approach is Hügelschäffer’s model (***Petrovic and Obradovic, 2010; Petrovic et al., 2011; Obradovic et al., 2013***).

The German engineer Fritz Hügelschäffer first proposed an oviform curve, shaped like an egg, by moving one of concentric circles along its *x*-axis constructing an asymmetric ellipse as reviewed elsewhere (***Schmidbauer, 1948; Ferréol, 2017***). A theoretical mathematical dependence for this curve was deduced elsewhere (***Petrovic and Obradovic, 2010; Petrovic et al., 2011***), which was later adapted by us in relation to the main measurements of the egg (i.e., its length, *L,* and maximum breadth, *B*) and carefully reviewed as applied to chicken eggs (***Narushin et al., 2020b***):

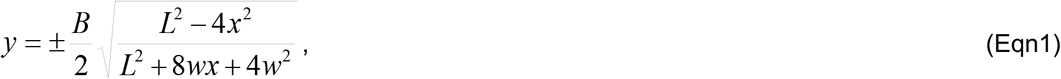

where *B* is the egg maximum breadth, *L* is the egg length, and *w* is the parameter that shows the distance between two vertical lines corresponding to the maximum breadth and the half length of the egg.

Hügelschäffer’s model works very well for three classical egg shapes, i.e., circular, elliptical and oval (***Figure 2A–D***). Indeed, when *L* = *B*, the shape becomes a circle and when *w* = 0 it becomes an ellipse. Therefore, the majority of avian egg shapes can be defined by the formula above (Eqn1). Unfortunately, Hügelschäffer’s model is not applicable in estimating the contours of pyriform eggs (***Figure 2E***). For instance, it is obvious even from visual inspection that the theoretical profile of the guillemot egg does not resemble its actual “real world” counterpart. Thus, Hügelschäffer’s model has some limitations in the description of the avian eggs, and one of those is a limited range of possible variations of the *w* value (***Narushin et al., 2020b***).

**Figure 2.**
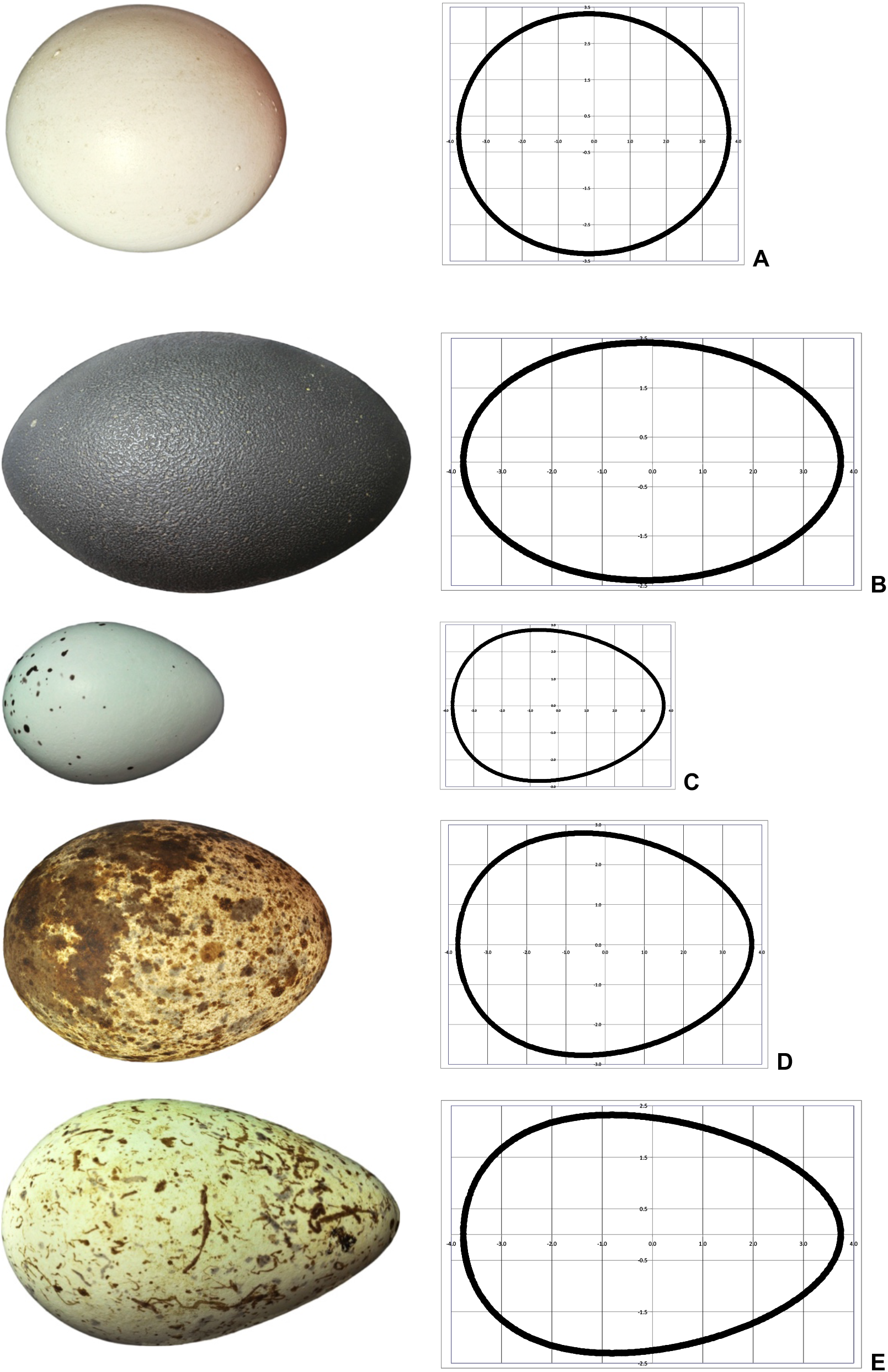
The images of eggs of the four main shapes from the following avian species:(**A**) ostrich, circular; (**B**) emu, elliptical; (**C**) song thrush, oval; (**D**) osprey, oval; and (**E**) guillemot pyriform; with their theoretical contours (on the right graphs) plotted using the Hügelschäffer’s model (Eqn1). The egg images were taken from Wikimedia Commons (Category: Eggs of the Natural History Collections of the Museum Wiesbaden), and their dimensions do not correspond to actual size due to scaling.

Based on the analysis of various formulae accumulated and available in the arsenal of egg geometry researchers (***Biggins et al., 2018***), one can admit a problem of a mathematical definition of pyriform (or conical) eggs to be the most difficult in comparison with all other egg shapes. With this in mind, the goal of this work was research aimed at developing a mathematical expression that would be able to accurately describe pyriform eggs and at devising a universal formula for avian eggs of any shape.

## Results

As a first step, we employed the data of numerous egg measurements represented by ***Romanoff and Romanoff*** (***1949***) for a standard hen’s egg, and produced the following formula for recalculation of *w* (see details in S1 Appendix):

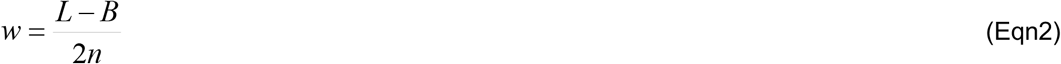

in which *n* is a positive number.

Inputting different numbers in Eqn2 and substituting the value of *w* into Eqn1, we can design different geometrical curves that resemble egg contours of other avian species (Figure 3).

**Figure 3.**
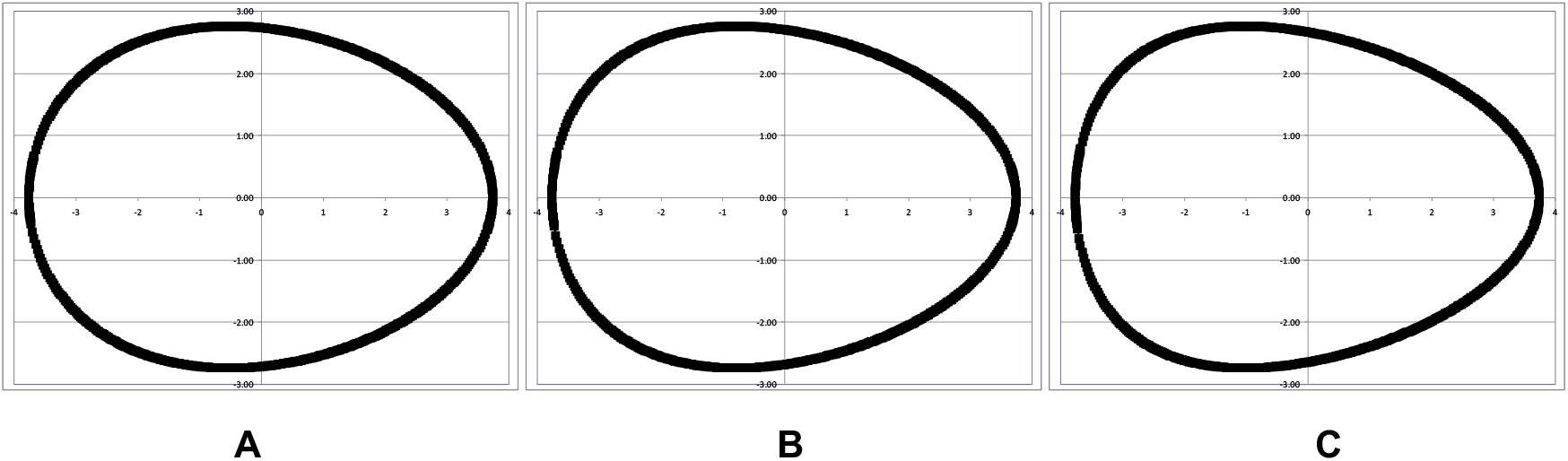
The egg contours plotted using Eqn1 and Eqn2 if: (**A**) *n* = 2, (**B**) *n* = 1.3, and (**C**) *n* = 1.

Thus, the principal limitation for Hügelschäffer’s model is the fact that *n* cannot be less than 1, which means that the maximum value of *w* is (*L*–*B*)/2. Otherwise, the obtained contour does not resemble the shape of any avian egg (Figure 4). This fact was investigated and well explained elsewhere (***Obradovic et al., 2013***).

**Figure 4.**
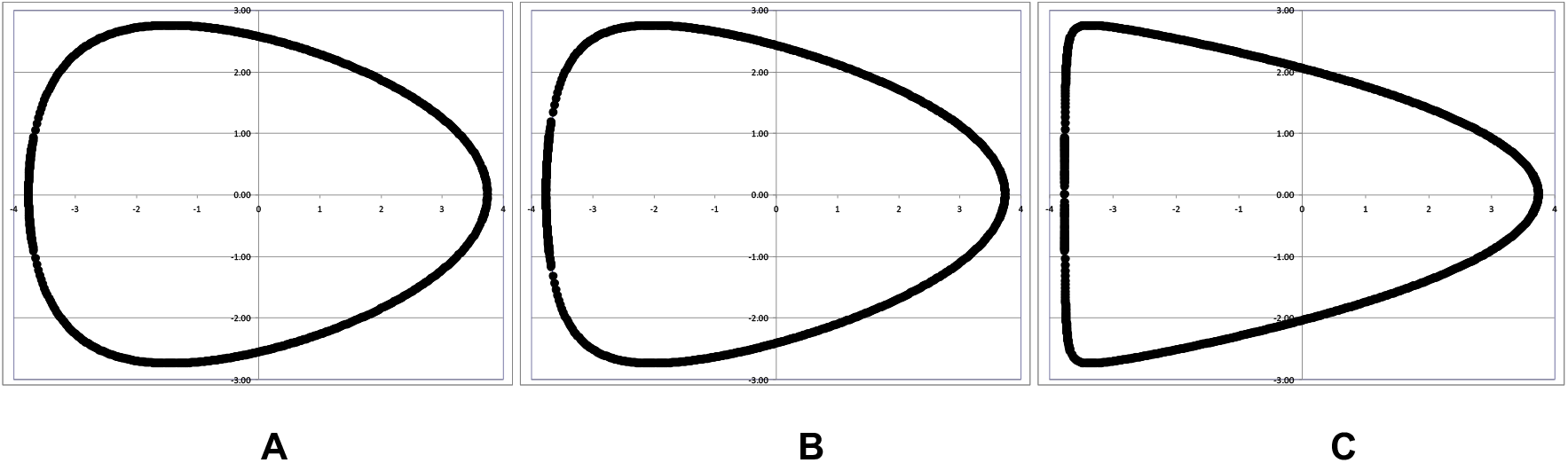
The egg contours plotted using Eqn1 and Eqn2 if: (**A**) *n* = 0.8, (**B**) *n* = 0.5, and (**C**) *n* = 0.3.

Such limitations explain why Hügelschäffer’s model cannot be used to describe the contours of pyriform eggs. The only way to make the shape of the pointed end of such eggs more conical is to use the *n* values less than 1, but in this case the obtained contours do not resemble any egg currently appearing in nature. In a series of mathematical computations, we deduced a formula for the pyriform egg shape (see details in S2 Appendix):

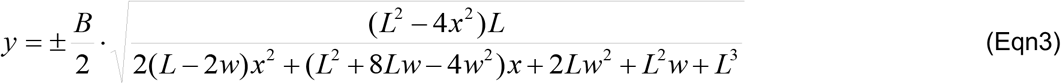

If we place the both contours, the pyriform Eqn3 and Hügelschäffer’s Eqn1 ones, together onto the same diagram (Figure 5), the presence of white area between them allows to arise a peculiar question: what to do with those eggs whose contours are tracing within this zone?

**Figure 5.**
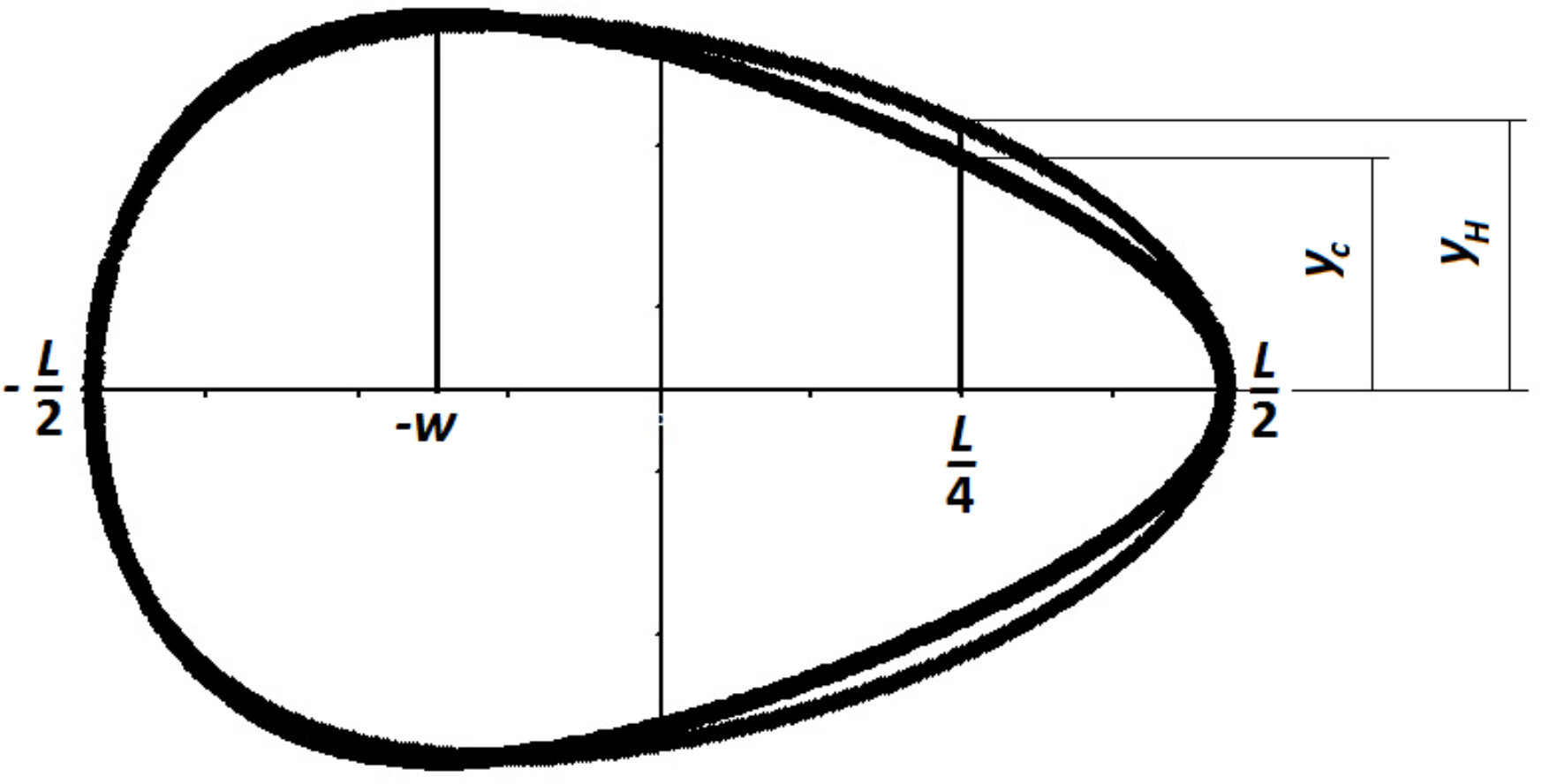
The contours of the egg plotted using the pyriform model according to Eqn3 (inner line) and the Hügelschäffer’s model according to Eqn1 (outer line).

If we choose any point on the *x*-axis within the interval [−*w…L*/2] corresponding to the white area between two models, there is obviously some difference, Δ*y*, between the values of the functions recalculated according to Hügelschäffer’s model, *y_H_* (Eqn1), and the pyriform one, *y_c_* (Eqn3), that tells how conical the egg is:

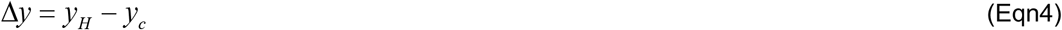

The subscript index ‘*c*’ was added only to designate that this function is related to its classic pyriform (conic) profile according to Eqn3 (*y_c_* does not differ from *y* in Eqn3). Maximum values of Δ*y* mean that the egg contour is related to its classic pyriform profile and can be expressed with Eqn3. When Δ*y* = 0, the egg shape has a classic ovoid profile (Hügelschäffer’s model) and is defined mathematically with Eqn1.

To fill this gap (Δ*y*) between the egg profiles according Eqn1 and Eqn3, the mathematical calculations were undertaken (S3 Appendix) being resulted in the final universal formula applicable for any avian egg:

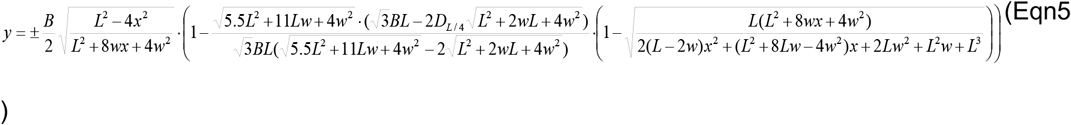

where *D*_*L*/4_ is egg diameter at the point of *L*/4 from the pointed end (Figure 5).

Both Eqn3 and Eqn5 were tested using pyriform eggs of different shape index (*SI*) and *w* to *L* ratio, and their validity were explicitly verified (Figure 6).

**Figure 6.**
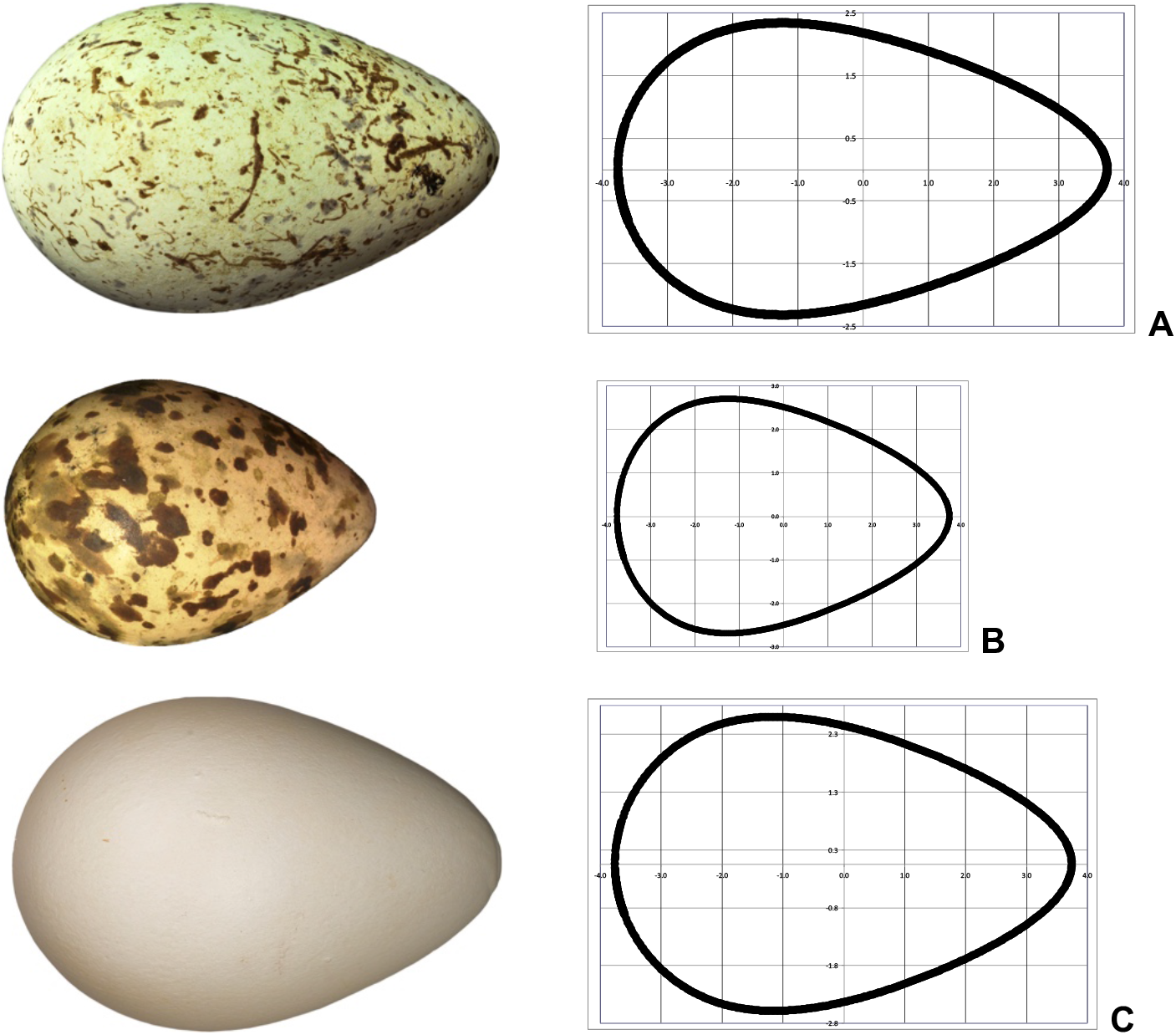
The images and their theoretical profiles of pyriform eggs of different shape index (*SI*) and *w* to *L* ratio: (**A**) a guillemot’s egg, *SI* = 0.58, *w/L* = 0.17; (**B**) a great snipe’s egg, *SI* = 0.69, *w/L* = 0.10; and (**C**) a king penguin’s egg, *SI* = 0.07, *w/L* = 1.8. The egg dimensions do not correspond to actual size due to scaling. The egg images were taken from Wikimedia Commons: (**A**) and (**B**) Category: Eggs of the Natural History Collections of the Museum Wiesbaden; and (**C**) Category: Bird eggs of the Muséum de Toulouse.

## Discussion

The common perception of “egg-shaped” is an oval, with a pointed end and a blunt end and the widest point nearest the blunt end, somewhat like a chicken’s egg. As we have demonstrated however, things can be far simpler (as in the case of the spherical eggs seen in owls, tinamous and bustards) or far more complicated (as in the case of pyriform eggs, e.g., seen in guillemots, waders and the two largest species of penguin). Evidence suggests (***Bradfield, 1951***) that egg shape is determined before the shell forms and by the underlying membranes. Why, in evolutionary terms, an egg is the shape that it is, is surprisingly under-studied. That is, although there are some previous investigations in the field of egg shape evolution (***Andersson, 1978; Stoddard et al., 2017, 2019; Birkhead et al., 2019***), we do not know how exactly this process occurred. In this context, it is the pyriform eggs (the ones that, in this study, we have incorporated in order to make the formula universal) that have attracted the most attention. In common sandpipers (and other waders) the pyriform shape is an adaptive trait ensuring that the (invariably) four eggs “fit together” in a nest (pointed ends innermost) to ensure maximum incubation surface against the mother’s brood patch (***Hewitson, 1831–1838***). In guillemots, the relative benefits of the pyriform shape to prevent eggs rolling off cliff edges have been much debated, however, to the best of our knowledge, this is far from certain (***Birkhead, 2016***). The selective advantage to being “oviform” rather than spherical is, according to ***Birkhead*** (***2016***), three-fold: First, given that a sphere has the smallest surface area to volume ratio of any geometric shape, there is a selective advantage to being roughly spherical as any deviation could lead to greater heat loss. Equally, non-spherical shapes are warmed more quickly and thus an egg may represent compromise morphology for most birds. A second consideration may well be, as in common sandpipers, related to “packaging” of the eggs in the brood, and the third could be related to the strength of the shell. In this final case, the considerations are that the egg needs to be strong enough so as not to rupture when sat on by the mother (a sphere is the best bet here), but weak enough to allow the chick to break out. As a compromise between to two, a somewhat elongated shape (be in elliptical, oval or pyriform) may represent a selective advantage.

In this study, we observed that applications of a mathematical apparatus in the area of oomorphology (***Mänd et al., 1986***) and egg shape geometry have developed from more simple formulae to more complex ones. In particular, the equation for the sphere would come first, being, then, modified into the equation for the ellipse by transforming the circle diameter into two unequal dimensions. Hügelschäffer’s model represented a mathematical approach to shift a vertical axis along the horizontal one. Finally, through the universal formula (Eqn5) we have provided here would allow to consider all possible egg profiles including the pyriform ones. For this, we would need only to measure the egg length, *L*, the maximum breadth, *B*, the distance *w* between the two vertical lines, corresponding to the maximum breadth and the half length of the egg, and the diameter, *D*_*L*/4,_ at the point of *L*/4 from the pointed end.

While we have provided evidence that our formula is universal for the overall shape of an avian egg, not every last contour of an avian egg may fit into the strict geometric framework of Eqn5. This is because natural objects are much more diverse and variable than mathematical objects. Nevertheless, generally speaking, we accept that the mountains are pyramidal, and the sun is round, although, in reality, their shapes only approximately resemble these geometric figures. In this regard, a methodological approach to assessing the shape of a particular bird egg would be to search for possible differences between the tested egg and its standard geometric shape (Eqn5). These distinctive criteria can (and should) be different for various purposes and specific research tasks. Perhaps, this would be the radius of the blunt and/or pointed end, or the skewness of one of the sections of the oval, or something else. The key message is that by introducing the universal egg shape formula we have expanded the arsenal of mathematics with another geometric figure that can safely be called a “real world” bird’s egg. The mathematical modelling of the egg shape and other egg parameters that we have presented here will be useful and important modus operandi for further stimulating the relevant theoretical and applied research in the fields of mathematics, engineering and biology (***Narushin et al., 2020a***).

In conclusion, a universal mathematical formula for egg shape has been proposed that is based on four parameters: egg length, maximum breadth, shift of the vertical axis and the diameter at one quarter of the egg length. This formula can theoretically describe any bird’s egg that exists in nature. Mathematical description of the sphere, the ellipsoid and the ovoid (all basic egg shapes) have already found numerous applications in a variety of academic disciplines including the biosciences, agriculture, architecture, aeronautics and mechanical engineering. We propose that this new formula will, similarly, have widespread application. We suggest that biological evolutionary processes such as egg formation are amenable to mathematical description, and may become the basis for the methodological concept of research in evolutionary biology.

In the course of the present analysis and search for the optimal mathematical approximation of oomorphology, i.e. the egg contours, we showed that our approach is as accurate as possible for the egg shape prediction. Based on the results of exploring the egg shape geometry models, we postulate here for the first time the theoretical formula that we have found as a universal equation for determining the contours of avian eggs. Our findings can be applied in a variety of fundamental and applied disciplines and serve as an impetus for the further development of scientific investigations using eggs as a research object.

## Materials and methods

To verify if the Hügelschäffer’s model (Eqn1) previously applied by us to chicken eggs (***Narushin et al., 2020b***) is valid to all possible egg shapes of various birds, we tested it on the following avian species: Ural owl (*Strix uralensis*) as a representative of circular eggs (Figure 2A), emu (*Dromaius novaehollandiae*) representing elliptical eggs (Figure 2B), song thrush (*Turdus philomelos*) and osprey (*Pandion haliaetus*) for oval eggs (Figures 2C and 2D), and guillemot (*Uria lomvia*) for pyriform eggs (Figure 2E).

In trying to establish if the novel formula of the pyriform contours (Eqn3) and the universal Eqn5 we developed here are valid for describing a variety of pyriform shapes, we applied them to the following avian species: guillemot (*Uria lomvia;* Figure 6A), great snipe (*Gallinago media;* Figure 6B), and king penguin (*Aptenodytes patagonicus;* Figure 6C).

For mathematical and standard statistical calculations, MS Excel and StatSoft programmes were exploited. As a part of our broader research project to develop more theoretical approaches for non-destructive evaluation of various characteristics of avian eggs (***Narushin et al., 2020a***), we did not handle eggs from wild birds or any valuable egg collection in this study. Where needed, we substituted actual eggs with their images and mathematical representational counterparts. To make it clear, we have considered a standard hen’s egg as represented by ***Romanoff and Romanoff*** (***1949***) and used their data of numerous egg measurements to deduce a formula for recalculation of *w* (S1 Appendix).

## Additional information

### Author contributions

All authors conceived and wrote the paper. Valeriy G Narushin performed the mathematical derivations and calculations.

### Additional files

S1 Appendix: Recalculation of w.

S2 Appendix: Mathematical description of pyriform eggs.

S3 Appendix: Inferring a universal formula for an avian egg.

### Data availability

All data generated or analysed during this study are included in the manuscript and supporting files.

## References

Andersson M. 1978. Optimal egg shape in waders. Ornis Fennica 55:105–109.

Baker DE. 2002. A geometric method for determining shape of bird eggs. Auk 119:1179–1186. doi: 10.1093/auk/119.4.1179.

Biggins JD, Thompson JE, Birkhead TR. 2018. Accurately quantifying the shape of birds’ eggs. Ecology and Evolution 8:9728–9738. doi: 10.1002/ece3.4412.

Birkhead TR. 2016. The Most Perfect Thing: Inside (and Outside) a Bird’s Egg. New York: Bloomsbury.

Birkhead TR, Thompson JE, Biggins JD, Montgomerie R. 2019. The evolution of egg shape in birds: selection during the incubation period. Ibis 161:605–618. doi: 10.1111/ibi.12658.

Bradfield JRG. 1951. Radiographic studies on formation of the hen’s egg shell. Journal of Experimental Biology 28:125–140.

Carter TC. 1968. The hen’s egg: A mathematical model with three parameters. British Poultry Science 9:165–171. doi: 10.1080/00071666808415706.

Cook JD. Equation to fit an egg. 2018 April 18 [cited 7 Aug 2020]. In: Blog [Internet]. John D. Cook Consulting. Available from: https://www.johndcook.com/blog/2018/04/18/equation-to-fit-an-egg/.

Ferréol R. Hügelschäffer egg. 2017 [cited 7 Aug 2020]. In: Encyclopédie des formes mathématiques remarquables. 2D Curves [Internet]. Robert Ferréol. Available from: http://www.mathcurve.com/courbes2d.gb/oeuf/oeuf.shtml.

Hewitson WC. 1831–1838. British Oology. Newcastle upon Tyne: Charles Empson.

Köller J. Egg curves and ovals. 2000 [cited 7 Aug 2020]. In: Mathematische Basteleien. More plane figures [Internet]. Jürgen Köller 1999–2020. Available from: http://www.mathematische-basteleien.de/eggcurves.htm.

Landa J. Rovnice vajíčka – jednoduchá jako Kolumbovo vejce. 2013 Mar 29 – Apr 23 [cited 7 Aug 2020]. In: Jirka Landa: Rovnice vajíčka [Internet]. Geoterra.EU 2008–. Available from: https://www.geoterra.eu/jirka-landa.

Mänd R, Nigul A, Sein E. 1986. Oomorphology: a new method. Auk 103:613–617. doi: 10.1093/auk/103.3.613.

Mytiai IS, Matsyura AV. 2017. Geometrical standards in shapes of avian eggs. Ukrainian Journal of Ecology 7:264–282. doi: 10.15421/2017_78.

Narushin VG, Lu G, Cugley J, Romanov MN, Griffin DK. 2020a. A 2-D imaging-assisted geometrical transformation method for non-destructive evaluation of the volume and surface area of avian eggs. Food Control 112:107112. doi: 10.1016/j.foodcont.2020.107112.

Narushin VG, Romanov MN, Lu G, Cugley J, Griffin DK. 2020b. Digital imaging assisted geometry of chicken eggs using Hügelschäffer’s model. Biosystems Engineering 197:45–55. doi: 10.1016/j.biosystemseng.2020.06.008.

Nishiyama Y. 2012. The mathematics of egg shape. International Journal of Pure and Applied Mathematics 78: 679–689.

Obradovic M, Malesevic B, Petrovic M, Djukanovic G. 2013. Generating curves of higher order using the generalisation of Hügelschäffer’s egg curve construction. Buletinul Ştiinţific al Universităţii,,Politehnica” din Timişoara: Seria Hidrotehnica 58:110–114.

Petrovic M, Obradovic M. 2010. The complement of the Hugelschaffer’s construction of the egg curve. In: Nestorović M, editor. 25th National and 2nd International Scientific Conference moNGeometrija 2010. Belgrade: Faculty of Architecture in Belgrade, Serbian Society for Geometry and Graphics. pp. 520–531.

Petrovic M, Obradovic M, Mijailovic R. 2011. Suitability analysis of Hugelschaffer’s egg curve application in architectural and structures’ geometry. Buletinul Instutului Politehnic din Iasi: Secţia Construcţii de maşini 57:115–122.

Pike TW. 2019. Quantifying the maculation of avian eggs using eggshell geometry. Ibis 161:686–693. doi: 10.1111/ibi.12708.

Preston FW. 1953. The shapes of birds’ eggs. Auk 70:160–182. doi: 10.2307/4081145.

Preston FW. 1968. The shapes of birds’ eggs: mathematical aspects. Auk 85;454–463. doi: 10.2307/4083294.

Romanoff AL, Romanoff AJ. 1949. The avian egg. New York: John Wiley & Sons Inc.

Schmidbauer H. 1948. Eine exakte Eierkurvenkonstruktion mit technischen Anwendungen. Elemente der Mathematik 3:67–68.

Smart IHM. 1991. Egg-shape in birds. In: Deeming DC, Ferguson MWJ, editors. Egg Incubation: Its Effects on Embryonic Development in Birds and Reptiles. Cambridge: Cambridge University Press. pp. 101–116. doi: 10.1017/CBO9780511585739.009.

Stoddard MC, Yong EH, Akkaynak D, Sheard C, Tobias JA, Mahadevan L. 2017. Avian egg shape: Form, function, and evolution. Science 356:1249–1254. doi: 10.1126/science.aaj1945.

Stoddard MC, Sheard C, Akkaynak D, Yong EH, Mahadevan L, Tobias JA. 2019. Evolution of avian egg shape: underlying mechanisms and the importance of taxonomic scale. Ibis 161:922–925. doi: 10.1111/ibi.12755.

Todd PH, Smart IHM. 1984. The shape of birds’ eggs. Journal of Theoretical Biology 106:239–243. doi: 10.1016/0022-5193(84)90021-3.

Troscianko J. 2014. A simple tool for calculating egg shape, volume and surface area from digital images. Ibis 156:874–878. doi: 10.1111/ibi.12177.

