## Appendices S1-S3 for "A universal formula for avian egg shape"

### Additional files

#### S1 Appendix: Recalculation of *w*

Let's consider a standard hen's egg as represented by ***Romanoff and Romanoff*** (***1949***) using their data of numerous egg measurements (***Figure S1***).

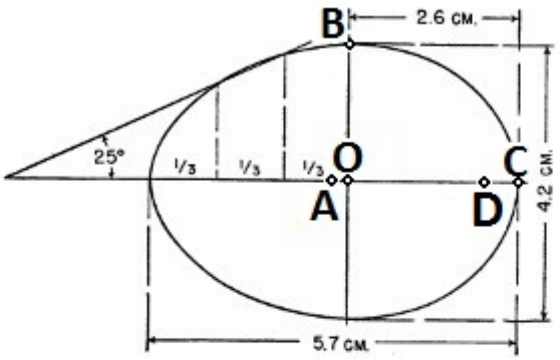


***Figure S1.*** A standard (or "ideal") chicken egg (1).

In this example, we selected the following five points on the egg image (***Figure S1***): A) a midpoint of the egg length, *L*/2; B) a point at the egg maximum breadth (*B)*; C) the utmost distant point of the blunt end of the egg; D) a point at a radius of an inner circle with the center in the O point, (where O is a cross point of the egg maximum breadth and length) so that the distances **OB** and **OD** are equal.

Then, we can state that **AO** = *w*, **OB** = **OD** = *B*/2, and

$$\mathbf{DC} = \frac{L}{2} - \frac{B}{2} - w \qquad \text{(EqnS1.1)}$$

Taking into consideration that:

$w = \frac{5.7}{2} - 2.6 = 0.25$ and **BC** = 2.6 – 2.1 = 0.5,

it is possible to conclude that **BC** = 2*w*. Then, EqnS1.1 can be rewritten as:

$$2w = \frac{L}{2} - \frac{B}{2} - w$$

$$w = \frac{L - B}{2 \cdot 3} \qquad \text{(EqnS1.2)}$$

For common usage of the Hügelschäffer's formula, we could rewrite EqnS1.2 as:

$$w = \frac{L - B}{2n} \qquad \text{(EqnS1.3)}$$

in which *n* is a positive number.

#### S1 Appendix References

**Romanoff AL**, Romanoff AJ. 1949. The avian egg. New York: John Wiley & Sons Inc.

**S2 Appendix: Mathematical description of pyriform eggs**

The image of the pointed end of the guillemot egg (***Figure 5E***) shows that it is complied with contours of a square parabola. If our assumption is correct, a blunt end of the pyriform egg will have a classic egg contour according to Hügelschäffer's model. The pointed end however should have a form of a parabola whose vertex lays on the *x*-axis, and the lines are rested against the extreme points of the egg maximum breadth (***Figure S2.1***).

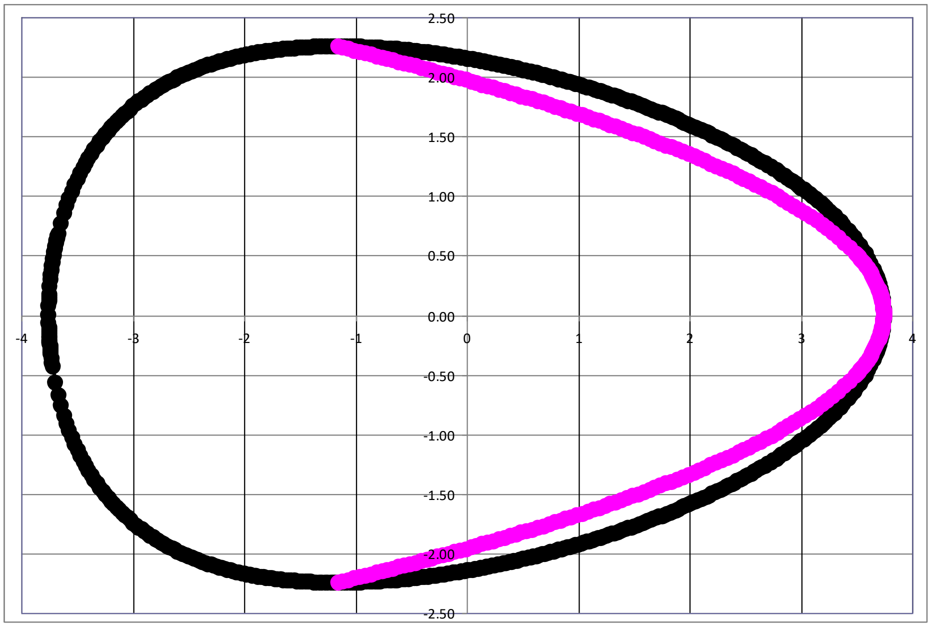


***Figure S2.1.*** Graphic representation of the guillemot egg profile using a parabola (pink line) and Hügelschäffer's model (black line).

Thus, the objective of our study was the theoretical definition of the pyriform egg with mathematical terms and plotted using both the Hügelschäffer's model and the parabola. Similar to ***Petrovic and Obradovic*** (***2010***), where principles of Newton's hyperbolism were used and the function $t(x)$

$$t(x) = 1 + \frac{2wx + w^2}{(L/2)^2} \qquad \text{(EqnS2.1)}$$

was added into the equation of the ellipse, obtaining a formula for a classic egg contour (or the Hügelschäffer's model):

$$\frac{x^2}{(L/2)^2} + \frac{y^2}{(B/2)^2} \cdot \left(1 + \frac{2wx + w^2}{(L/2)^2}\right) = 1 \qquad \text{(EqnS2.2)}$$

we assumed to follow the same principle for a pyriform profile, so another function that we conditionally defined as $p(x)$ and called a 'pyriform function' was used in the EqnS2.2 instead of $t(x)$:

$$\frac{x^2}{(L/2)^2} + \frac{y^2}{(B/2)^2} \cdot p(x) = 1 \qquad \text{(EqnS2.3)}$$

Wherefrom:

$$p(x) = \frac{B^2}{4} \cdot \frac{L^2 - 4x^2}{y^2 L^2} \qquad \text{(EqnS2.4)}$$

According to our assumption, a vertex of the parabola that corresponds to the pointed end of the classic egg contour (Hügelschäffer's model) lays on the $x$-axis at $x = L/2$ and the lines are rested against the extreme points of the egg maximum breadth, $B$, which is shifted from the $y$-axis at the value of $x = -w$, while the blunt end has the form of Hügelschäffer's model (***Figure S2.2***).

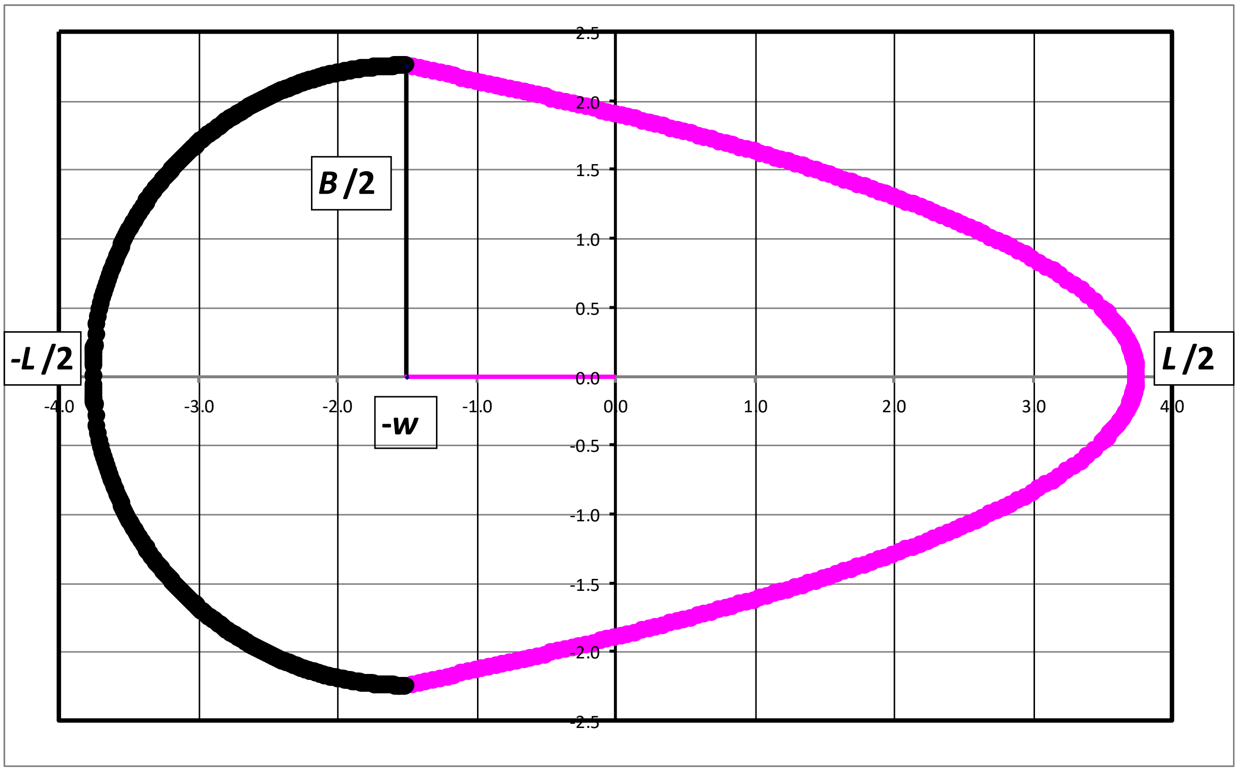


***Figure S2.2.*** Geometry of the pyriform egg.

Taking into account that a vertex of a horizontal parabola is located on the *x*-axis, thus grounding upon theoretical principles for the following function as reviewed elsewhere (***Cognero, 2019***) and the data which follows from ***Figure S2.2***, when $y = 0$, $x = L/2$; and when $x = -w$, $y = B/2$, we obtained the functional dependence of the parabolic end of the pyriform egg contour:

$$y_p = \pm\frac{B}{2}\cdot\sqrt{\frac{L-2x_p}{L+2w}} \qquad \text{(EqnS2.5)}$$

Considering that the pyriform egg contour consists of two geometrical figures, the parabola and Hügelschäffer's model, the values *x* and *y* of the pointed and blunt ends are different, so the subscript index '*p*' would correspond to the meanings of *x* and *y* of the parabolic part, and the ones for the Hügelschäffer's part would have the subscript index '*H*'. Thus, the interval for $x_p$ is [$-w$, $L/2$] and the one for $x_H$ is [$-L/2$, $-w$].

Inputting EqnS2.5 into EqnS2.4 we inferred the pyriform function $p(x_p)$ for the parabolic part of the egg:

$$p(x_p) = \frac{(L + 2x_p)(L + 2w)}{L^2} \qquad \text{(EqnS2.6)}$$

Then, the pyriform function $p(x_H)$ for Hügelschäffer's part of the egg is expressed accordingly from EqnS2.1 thus:

$$p(x_H) = \frac{L^2 + 8wx_H + 4w^2}{L^2} \qquad \text{(EqnS2.7)}$$

**To produce a universal formula for recalculating the pyriform multiplier $p(x)$, we would need to unite the both equations for $p(x_p)$ and $p(x_H)$ into one.** For this, each EqnS2.6 and EqnS2.7 were transformed as follows:

$$p(x_p) = \left(1 + 2 \cdot \frac{x_p}{L}\right) \cdot \left(1 + 2 \cdot \frac{w}{L}\right) \qquad \text{(EqnS2.8)}$$

$$p(x_H) = 1 + 8 \cdot \frac{w}{L} \cdot \frac{x_H}{L} + 4 \cdot \left(\frac{w}{L}\right)^2 \qquad \text{(EqnS2.9)}$$

If we express the values of $x$ in terms of the egg length $L$ as $x = aL$, wherefrom $a = x/L$, then, for the parabolic part of the egg the values of $a_p$ would be within the interval $a_p = [-w/L;\ 1/2]$, and for the Hügelschäffer's one within $a_H = [-1/2;\ -w/L]$ (***Figure S2.2***), so EqnS2.8 and EqnS2.9 were rewritten as follows:

$$p(x_p) = (1 + 2a_p) \cdot \left(1 + 2 \cdot \frac{w}{L}\right) \quad \text{(EqnS2.10)}$$

$$p(x_H) = 1 + 8a_H \cdot \frac{w}{L} + 4 \cdot \left(\frac{w}{L}\right)^2 \quad \text{(EqnS2.11)}$$

The united function *p*(*x*) should satisfy the both EqnS2.10 and EqnS2.11 within the whole interval of *a* = [–1/2; 1/2]. Prior to start an approximation procedure of such combination, we defined possible variations of the meanings of *w*/*L*. In accords with the Hügelschäffer's model, *w* cannot be less than 0. In this case, the egg image just revolves for 180° over the *x*-axis and, thus, the minimum possible value for *w* is 0. When *w* = 0, the classic egg contour transforms into the ellipse (***Petrovic and Obradovic***, ***2010***). As above mentioned, the maximum possible meaning of *w* is (*L*–*B*)/2. Thus, the minimum possible value of *w*/*L* is 0, and the maximum one is:

$$\frac{w_{\max}}{L} = \frac{L - B}{2L} = \frac{1 - SI}{2} \quad \text{(EqnS2.12)}$$

where *SI* is the egg shape index (maximum breadth to length ratio). Then, the maximum value of *w* would be when the value of *SI* is minimal. We found that the mostly elongated eggs, i.e., with the least shape index (0.55 to 0.57), are inherent in the maleos (*Macrocephalon maleo*) and long-tailed sylph (*Aglaiocercus kingii*) (***Stoddard et al., 2017***; ***Hauber, 2014***). Nevertheless, the similar approach to describing oomorpholgy can be used not only for avian species. For example, the eggs of American crocodiles and some dinosaurs (***Wiemann et al., 2018***) have the shape index values close to 0.5, so we assumed this as a minimal value for *SI*. Then, as follows from EqnS2.12, $w_{\max}/L$ = 0.25, and performing the approximation procedure, we should deal with the interval of *w*/*L* = [0; 0.25], dividing it into five equal subintervals of the length 0.05 with the following subinterval endpoints: 0, 0.05, 0.1, 0.15, 0.2 and 0.25.

The meanings of $p(x_H)$ and $p(x_p)$ (EqnS2.10 and EqnS2.11) were recalculated by inputting the correspondent values of *a* for the Hügelschäffer's part of the egg, $a_H$ = [–1/2; –*w*/*L*], and for the parabolic one, [–*w*/*L*; 1/2]. As a result, the diagram in ***Figure S2.3*** was plotted.

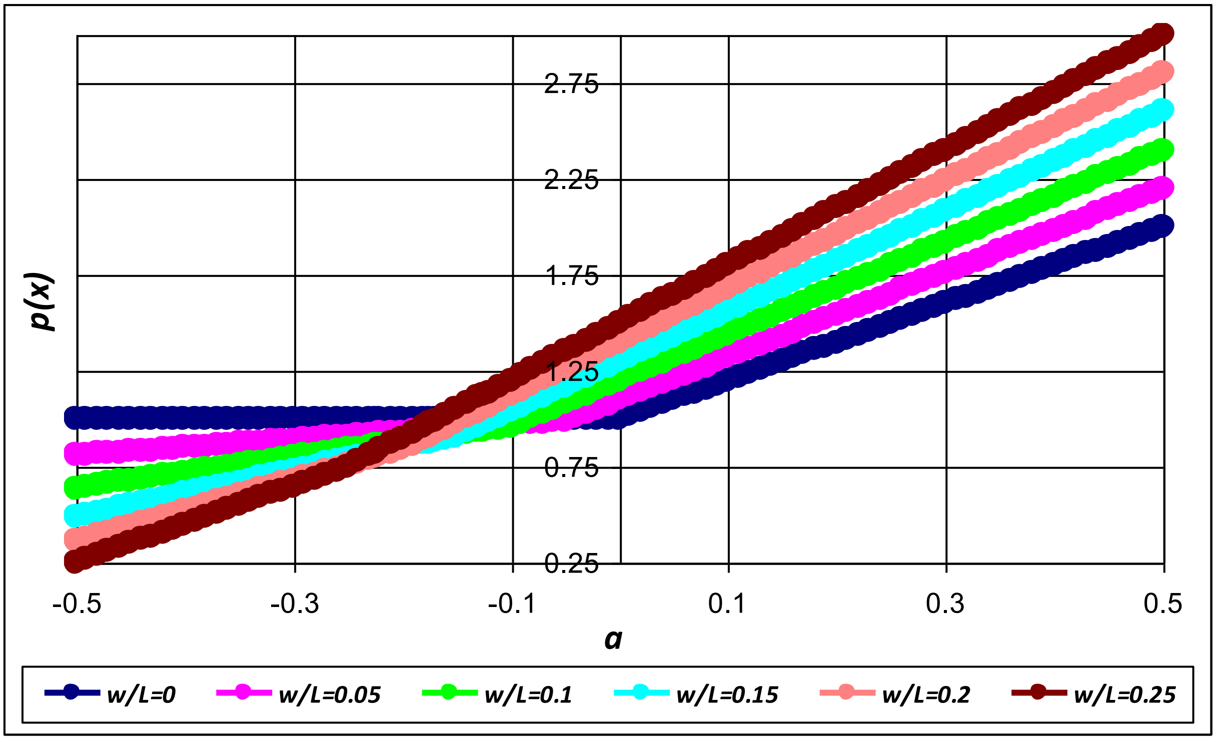


***Figure S2.3.*** The curves of *p*(*x*) as a function of *a* according to EqnS2.10 and EqnS2.11 for the values of *w*/*L* ranged between 0 and 0.25.

The functions of *p*(*x*) in ***Figure S2.3*** are related to piecewise functions (***Egbert, 2016***), so we can unite these into one using a numerical procedure.

At first, we intended to define an approximating function that would double the combination of the obtained curves (***Figure S2.3***) as accurate as possible. Taking into account that the curves consist of two linear functions (EqnS2.10 and EqnS2.11), principles of linear algebra (***Lay et al., 2016***) were used for this approximation. The most accurate results were obtained using the following formula:

$$p(x) = \frac{c_1 + c_2 a + c_3 a^2}{c_4 + c_5 a + a^2} \qquad \text{(EqnS2.13)}$$

in which $c_1 \ldots c_5$ are constant coefficients to be defined.

The results of the approximation with EqnS2.13 are shown in ***Figure S2.4***.

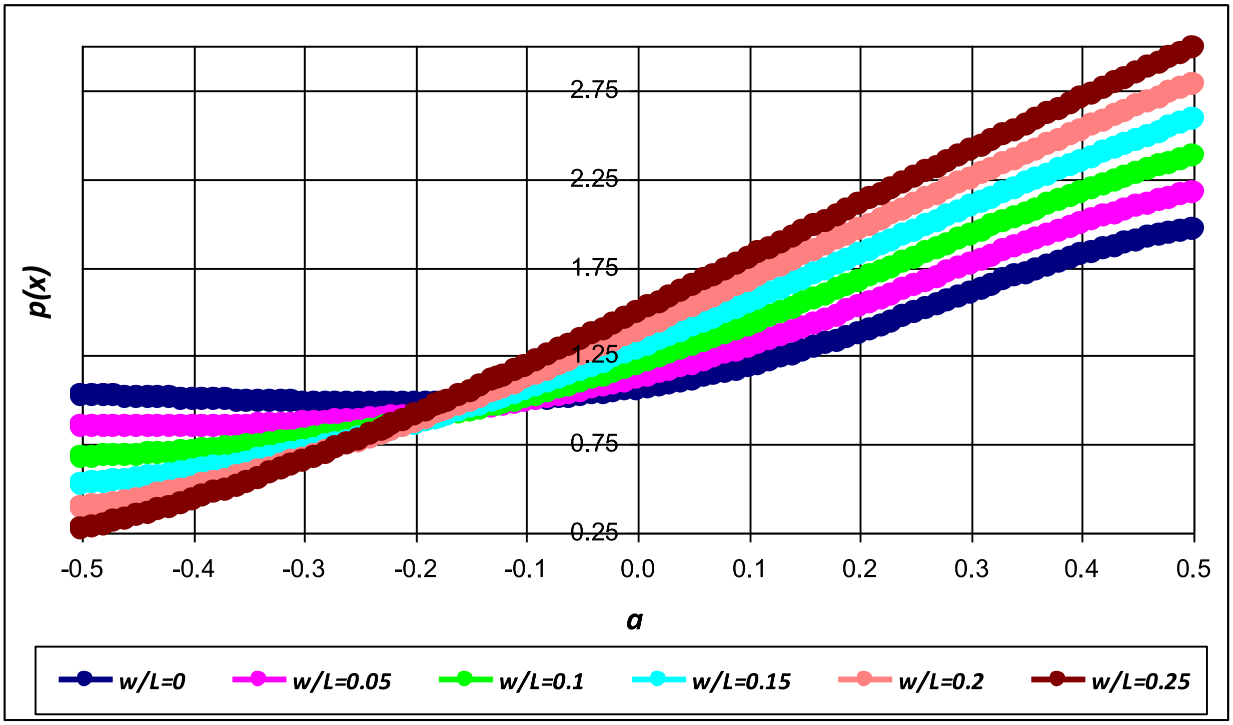


***Figure S2.4.*** The approximated curves $p(x)$ using EqnS2.13.

The obtained coefficients for each meaning of $w/L$ are presented in ***Table S2.1*** together with the coefficient of determination ($R^2$) between the results calculated using EqnS2.13 and EqnS2.10 or EqnS2.11, respectively.

***Table S2.1.*** Meanings of the coefficients $c_1 \ldots c_5$ in EqnS2.13.

| ***w*/*L*** | $c_1$ | $c_2$ | $c_3$ | $c_4$ | $c_5$ | $R^2$ |
|---|---|---|---|---|---|---|
| 0 | 0.316 | -0.320 | 1.493 | 0.297 | -0.556 | 0.9962 |
| 0.05 | 0.517 | 0.024 | 1.415 | 0.461 | -0.609 | 0.9979 |
| 0.10 | 0.851 | 0.814 | 1.548 | 0.711 | -0.537 | 0.9988 |
| 0.15 | 1.447 | 2.484 | 2.286 | 1.124 | -0.221 | 0.9994 |
| 0.20 | 2.617 | 6.128 | 4.612 | 1.886 | 0.635 | 0.9997 |
| 0.25 | 5.171 | 14.714 | 11.246 | 3.467 | 2.828 | 0.9999 |

To make EqnS2.13 valid for all meanings of *w*/*L* (***Figure S2.4***), each coefficient *c* was defined as a function *c* = *f*(*w*/*L*) and, then, approximated with relevant equations (***Figure S2.5***).

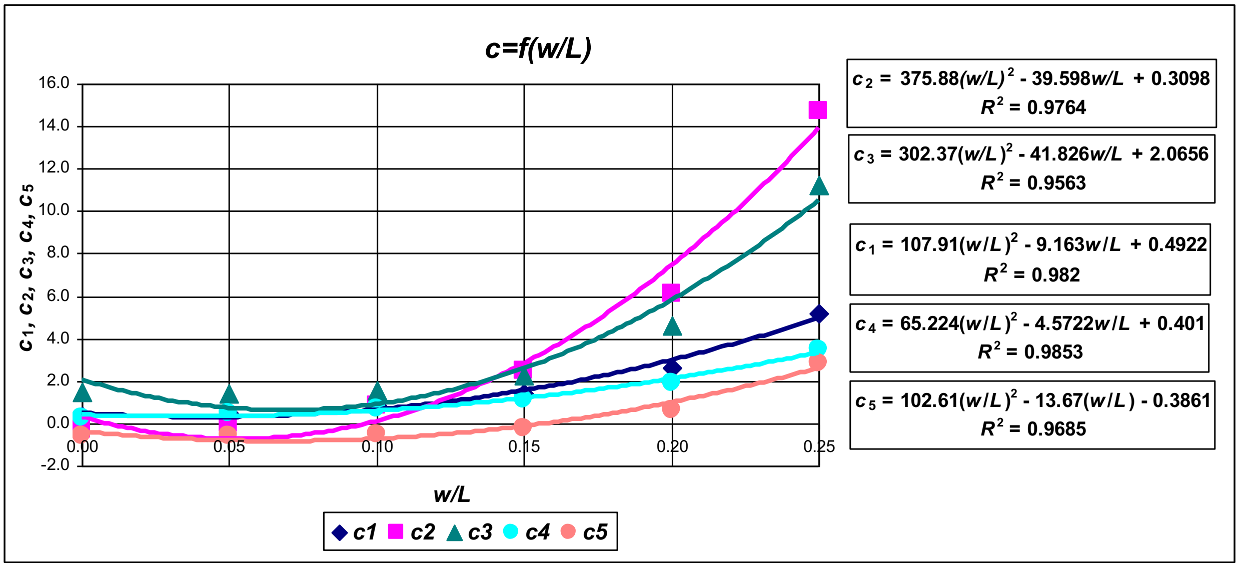


***Figure S2.5.*** The approximation of the coefficients $c_1 \ldots c_5$ with square polynomials.

As a result, the final approximating formula was defined as follows:

$$p(x)=\frac{107.91\cdot\left(\frac{w}{L}\right)^2-9.16\cdot\frac{w}{L}+0.49+\left(375.88\cdot\left(\frac{w}{L}\right)^2-39.6\cdot\frac{w}{L}+0.31\right)\cdot\frac{x}{L}+\left(302.37\cdot\left(\frac{w}{L}\right)^2-41.83\cdot\frac{w}{L}+2.07\right)\cdot\left(\frac{x}{L}\right)^2}{65.22\cdot\left(\frac{w}{L}\right)^2-4.57\cdot\frac{w}{L}+0.4+\left(102.61\cdot\left(\frac{w}{L}\right)^2-13.67\cdot\frac{w}{L}-0.39\right)\cdot\frac{x}{L}+\left(\frac{x}{L}\right)^2}$$

(EqnS2.14)

Considering EqnS2.3, the function that defines the contours of the pyriform eggs can be represented as:

$$y=\pm\frac{B}{2}\cdot\sqrt{\frac{L^2-4x^2}{L}}\cdot\sqrt{\frac{65.22\cdot\left(\frac{w}{L}\right)^2-4.57\cdot\frac{w}{L}+0.4+\left(102.61\cdot\left(\frac{w}{L}\right)^2-13.67\cdot\frac{w}{L}-0.39\right)\cdot\frac{x}{L}+\left(\frac{x}{L}\right)^2}{107.91\cdot\left(\frac{w}{L}\right)^2-9.16\cdot\frac{w}{L}+0.49+\left(375.88\cdot\left(\frac{w}{L}\right)^2-39.6\cdot\frac{w}{L}+0.31\right)\cdot\frac{x}{L}+\left(302.37\cdot\left(\frac{w}{L}\right)^2-41.83\cdot\frac{w}{L}+2.07\right)\cdot\left(\frac{x}{L}\right)^2}}$$

(EqnS2.15)

Judging from the appropriate graphic (***Figure S2.6***) and further theoretical analysis of the obtained equation EqnS2.15, it showed a considerable drawback: at the point of $x = -w$, $y$ should be equal to $B/2$.

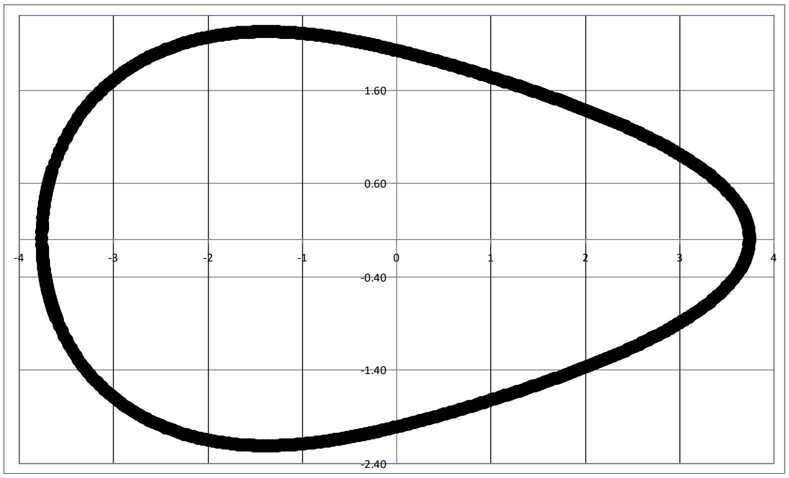


***Figure S2.6.*** The contour of an actual guillemot egg plotted using EqnS2.15.

This principle was thoroughly described in our previous paper (***Narushin et al., 2020***). Apart from looking very similar to the actual guillemot egg contour, the maximum value of $y$ that is supposed to be equal to $B/2$ = 2.25 cm in our case, equals to 2.21 cm. This discrepancy would require a further detailed revisit of EqnS2.15 as follows:

if $x = -w$, $y_{max} = B/2$, and considering EqnS2.15, that is valid when

$$\sqrt{\frac{L^2-4w^2}{L}}\cdot\sqrt{\frac{65.22\cdot\left(\frac{w}{L}\right)^2-4.57\cdot\frac{w}{L}+0.4-\left(102.61\cdot\left(\frac{w}{L}\right)^2-13.67\cdot\frac{w}{L}-0.39\right)\cdot\frac{w}{L}+\left(\frac{w}{L}\right)^2}{107.91\cdot\left(\frac{w}{L}\right)^2-9.16\cdot\frac{w}{L}+0.49-\left(375.88\cdot\left(\frac{w}{L}\right)^2-39.6\cdot\frac{w}{L}+0.31\right)\cdot\frac{w}{L}+\left(302.37\cdot\left(\frac{w}{L}\right)^2-41.83\cdot\frac{w}{L}+2.07\right)\cdot\left(\frac{w}{L}\right)^2}}=1$$

(EqnS2.16)

The obtained equation EqnS2.16 demonstrated a bias and was not true for the whole possible interval of $w/L$ meanings. Numerical methods in adjusting EqnS2.16 enabled us to stipulate that the main error occurs when the results of the approximated calculations do not coincide with the initial ones at three basic points $p(-L/2)$, $p(-w)$ and $p(L/2)$, or when $a = -1/2$, $a = -w$ and $a = 1/2$. Thus, to circumvent this obstacle, we undertook another approach. For each meaning of $w/L$ chosen previously, we determined the meanings at three basic points (***Table S2.2***), which were recalculated from EqnS2.10 and EqnS2.11 accordingly.

***Table S2.2.*** The meanings of $a$ and $p(x)$ at three basic points $p(-L/2)$, $p(-w)$ and $p(L/2)$.

| ***w/L*** | ***a* = –1/2** | ***a* = –*w*** | ***a* = 1/2** | ***p*(–*L*/2)** | ***p*(–*w*)** | ***p*(*L*/2)** |
|---|---|---|---|---|---|---|
| 0 | –0.5 | 0.00 | 0.5 | 1.00 | 1.00 | 2.00 |
| 0.05 | –0.5 | –0.05 | 0.5 | 0.81 | 0.99 | 2.20 |
| 0.10 | –0.5 | –0.1 | 0.5 | 0.64 | 0.96 | 2.40 |
| 0.15 | –0.5 | –0.15 | 0.5 | 0.49 | 0.91 | 2.60 |
| 0.20 | –0.5 | –0.2 | 0.5 | 0.36 | 0.84 | 2.80 |

| | | | | | | |
|---|---|---|---|---|---|---|
| 0.25 | 5.171 | 14.714 | 11.246 | 3.467 | 2.828 | 0.9999 |

The results of the approximation of the data from ***Table S2.2*** are given graphically in ***Figure S2.7***.

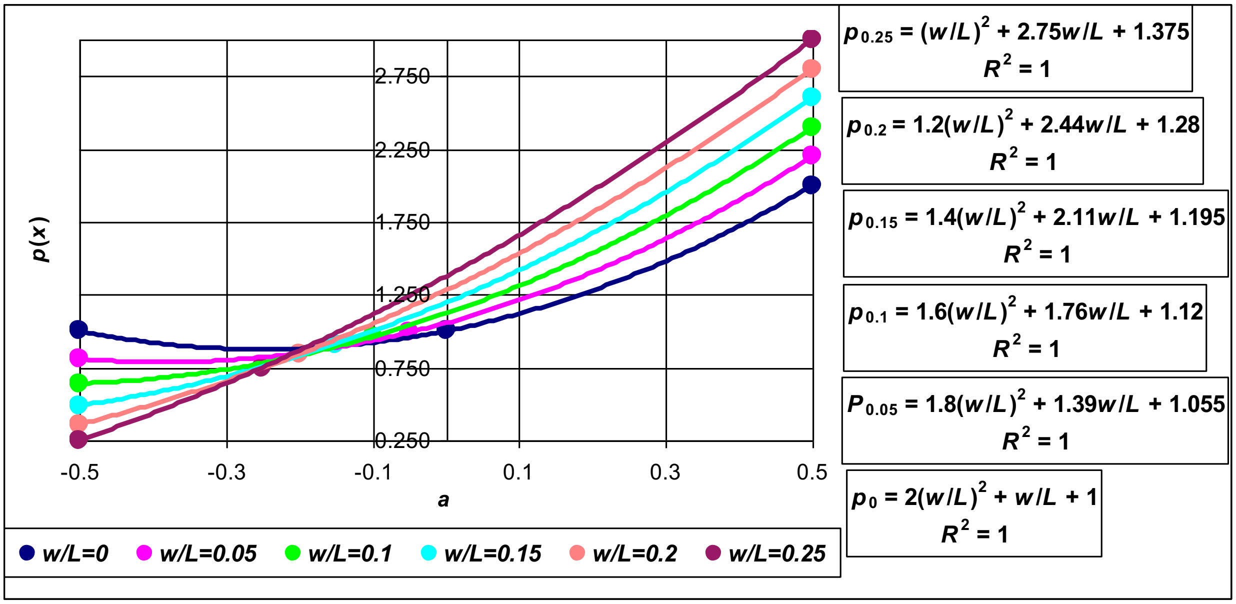


***Figure S2.7***. The results of the approximation of the data from ***Table S2.2***.

All the curves showed very accurate results expressed with square polynomials. Similar to the previous approximating approach, to make the results valid for all meanings of *w*/*L* (***Figure S2.7***), each coefficient *c* was defined as a function *c* = *f*(*w*/*L*), where coefficients $c_1 \dots c_3$ correspondingly fit the following condition:

$$p_{w/L} = c_1 \cdot \left(\frac{w}{L}\right)^2 + c_2 \cdot \frac{w}{L} + c_3 \qquad \text{(EqnS2.17)}$$

in which $p_{w/L}$ means the value of *p*(*x*) (EqnS2.10 and EqnS2.11) for the respective value of *w*/*L*.

The results of the coefficients approximations are shown graphically in ***Figure S2.8***.

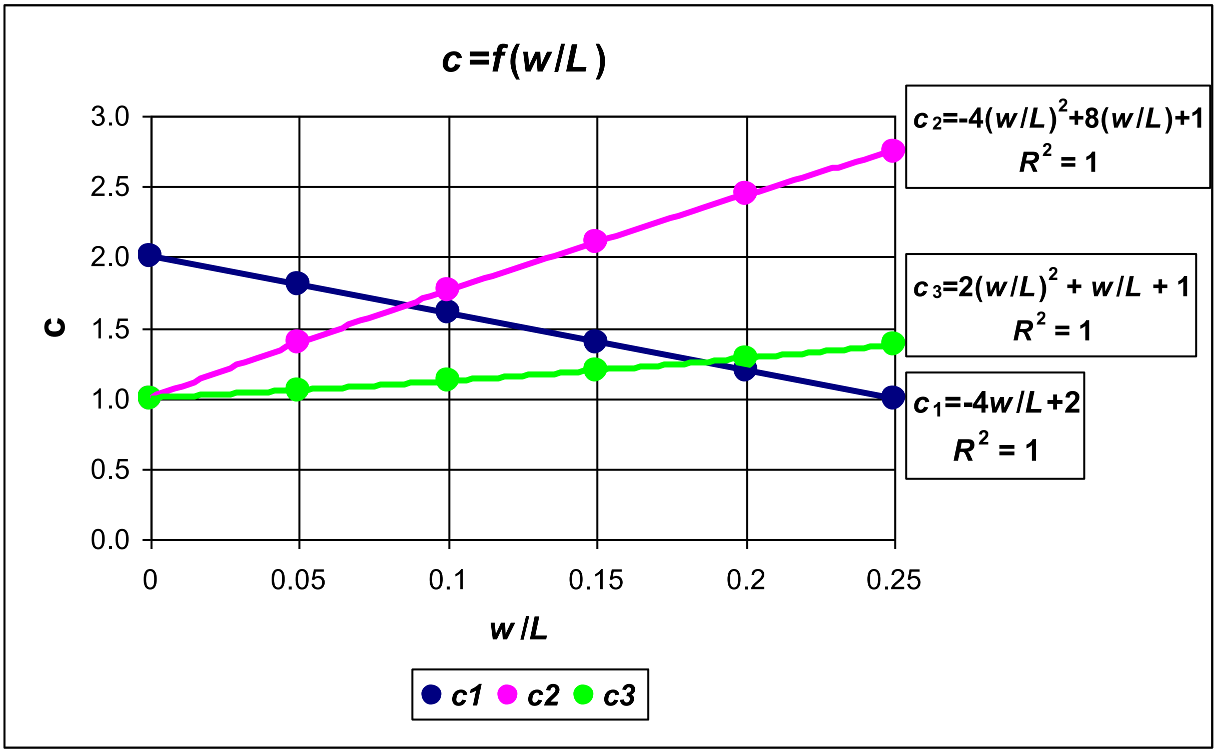


***Figure S2.8.*** The results of the approximation of the coefficients $c_1 \ldots c_3$ in the equations of *Figure S2.5.*

Then, the penultimate approximating formula was defined as follows:

$$p(x) = \frac{2(L-2w)x^2 + (L^2 + 8Lw - 4w^2)x + 2Lw^2 + L^2w + L^3}{L^3} \qquad \text{(EqnS2.18)}$$

And finally, the function that should satisfy the geometrical description of the pyriform eggs can be inferred after substituting EqnS2.18 into EqnS2.3 as follows:

$$y = \pm\frac{B}{2} \cdot \sqrt{\frac{(L^2 - 4x^2)L}{2(L-2w)x^2 + (L^2 + 8Lw - 4w^2)x + 2Lw^2 + L^2w + L^3}} \qquad \text{(EqnS2.19)}$$

The undertaken test of substituting $x = -w$ into EqnS2.19 demonstrated that the meaning of $y_{max} = \pm B/2$, and EqnS2.19 fully satisfies the basic condition for the universal geometrical description of avian eggs:

$$\sqrt{\frac{(L^2-4w^2)L}{2(L-2w)w^2+(L^2+8Lw-4w^2)w+2Lw^2+L^2w+L^3}}=1$$

$$L^3-4Lw^2-2Lw^2+4w^3+L^2w+8Lw^2-4w^3-2Lw^2-L^2w-L^3=0$$

The above iterations were resulted in the following basic formulae:

$$p(x)=\frac{2(L-2w)x^2+(L^2+8Lw-4w^2)x+2Lw^2+L^2w+L^3}{L^3} \qquad \text{(EqnS2.20)}$$

and

$$y=\pm\frac{B}{2}\cdot\sqrt{\frac{(L^2-4x^2)L}{2(L-2w)x^2+(L^2+8Lw-4w^2)x+2Lw^2+L^2w+L^3}} \qquad \text{(EqnS2.21)}$$

A graphical representation of EqnS2.21 (or Eqn3) for the images of typical representatives of the pyriform eggs of different variations in the values of their shape index and $w/L$ ratio also showed its validity (***Figure 6***).

#### S2 Appendix References

**Cognero**. 7–1 Parabolas. 2019 [cited 7 Aug 2020]. In: eSolutions Manual – Powered by Cognero [Internet]. North Ridgeville: North Ridgeville City Schools 2020. Available from: https://www.nrcs.net/Downloads/T7-1%20hw5.pdf.

**Egbert N**. Piecewise functions. 2016 Fall [cited 7 Aug 2020]. In: Teaching. Fall 2016: MA 158 [Internet]. West Lafayette: Nicholas Egbert, Mathematics Department, Purdue University 2018– . Available from: https://www.math.purdue.edu/~egbertn/fa2016/notes/lesson9.pdf.

**Hauber ME**. 2014. The Book of Eggs: a Life-size Guide to the Eggs of Six Hundred of the World's Bird Species. Chicago: University of Chicago Press.

**Lay DC**, Lay SR, McDonald JJ. 2016. Linear algebra and its applications. 5th ed. Boston: Pearson.

**Narushin VG**, Romanov MN, Lu G, Cugley J, Griffin DK. 2020. Digital imaging assisted geometry of chicken eggs using Hügelschäffer's model. *Biosystems Engineering* **197**:45–55. doi: 10.1016/j.biosystemseng.2020.06.008.

**Petrovic M**, Obradovic M. 2010. The complement of the Hugelschaffer's construction of the egg curve. In: Nestorović M, editor. 25th National and 2nd International Scientific Conference moNGeometrija 2010. Belgrade: Faculty of Architecture in Belgrade, Serbian Society for Geometry and Graphics. pp. 520–531.

**Stoddard MC**, Yong EH, Akkaynak D, Sheard C, Tobias JA, Mahadevan L. 2017. Avian egg shape: Form, function, and evolution. *Science* **356**:1249–1254. doi: 10.1126/science.aaj1945.

**Wiemann J**, Yang TR, Norell MA. 2018. Dinosaur egg colour had a single evolutionary origin. *Nature* **563**:555–558. doi: 10.1038/s41586-018-0646-5.

#### S3 Appendix: Inferring a universal formula for an avian egg

For enabling us to express the contours between these two classic models, the pyriform one (EqnS2.21, or Eqn3) and the ovoid one (Eqn1), we assumed that the pyriform function $p(x)$ (EqnS2.20) can be represented as a product of the Hügelschäffer's function, $t(x)$, (EqnS2.1) and some multiplier, $M_p$, that we coined as a *pyriform multiplier*:

$$p(x) = t(x) \cdot M_p \qquad \text{(EqnS3.1)}$$

Then, as follows from (EqnS2.3)

$$y_H = \frac{B}{2L} \cdot \sqrt{\frac{L^2 - 4x^2}{p(x)}} \qquad \text{(EqnS3.2)}$$

the formula for the conic shapes can be expressed accounting EqnS3.1 as follows:

$$y_c = \frac{B}{2L} \cdot \sqrt{\frac{L^2 - 4x^2}{t(x) \cdot M_p}} \qquad \text{(EqnS3.3)}$$

wherefrom considering (Eqn4):

$$\Delta y = y_H \cdot \left(1 - \frac{1}{\sqrt{M_p}}\right) \qquad \text{(EqnS3.4)}$$

As follows from EqnS2.20 and EqnS2.1, the pyriform multiplier is defined as follows:

$$M_p = \frac{2(L-2w)x^2 + (L^2 + 8Lw - 4w^2)x + 2Lw^2 + L^2 w + L^3}{L(L^2 + 8wx + 4w^2)} \quad \text{(EqnS3.5)}$$

Then, considering EqnS3.4:

$$\Delta y = y_H \cdot \left(1 - \sqrt{\frac{L(L^2 + 8wx + 4w^2)}{2(L-2w)x^2 + (L^2 + 8Lw - 4w^2)x + 2Lw^2 + L^2 w + L^3}}\right) \quad \text{(EqnS3.6)}$$

As follows from Eqn4 and EqnS3.6, the classic pyriform (conic) shape can be expressed with the following equation:

$$y_c = y_H \sqrt{\frac{L(L^2 + 8wx + 4w^2)}{2(L-2w)x^2 + (L^2 + 8Lw - 4w^2)x + 2Lw^2 + L^2 w + L^3}} \quad \text{(EqnS3.7)}$$

If to substitute the formula for $y_H$ (Eqn1), as expected EqnS3.7 is identical to EqnS2.21 (or Eqn3).

If an egg profile is located just between Hügelschäffer's and pyriform curves, we would need in this case to take only a respective part of $\Delta y$, which can be defined as some coefficient $k$. Then, if we express a mathematical function of such eggs as $y_{H/c}$:

$$y_{H/c} = y_H - k \cdot \Delta y \quad \text{(EqnS3.8)}$$

or

$$y_{H/c} = y_H \left(1 - k\left(1 - \sqrt{\frac{L(L^2 + 8wx + 4w^2)}{2(L-2w)x^2 + (L^2 + 8Lw - 4w^2)x + 2Lw^2 + L^2 w + L^3}}\right)\right)$$

(EqnS3.9)

The meanings of $k$ are within the interval [0…1]. When $k = 0$, the egg shape is expressed with the Hügelschäffer's model. If $k = 1$, then, it appears to be of classic pyriform (conic) one, so (EqnS3.9) can be considered as **the universal formula for any avian egg.**

**Independent validation of the universal formula**

To check if (EqnS3.9) is valid, let us assume that $k = 0.5$. The graphic representations of the egg contours (***Figure S3***) with $k$ = 0, 0.5 and 1 suggested that (EqnS3.9) can be used for practical calculations of these egg profiles.

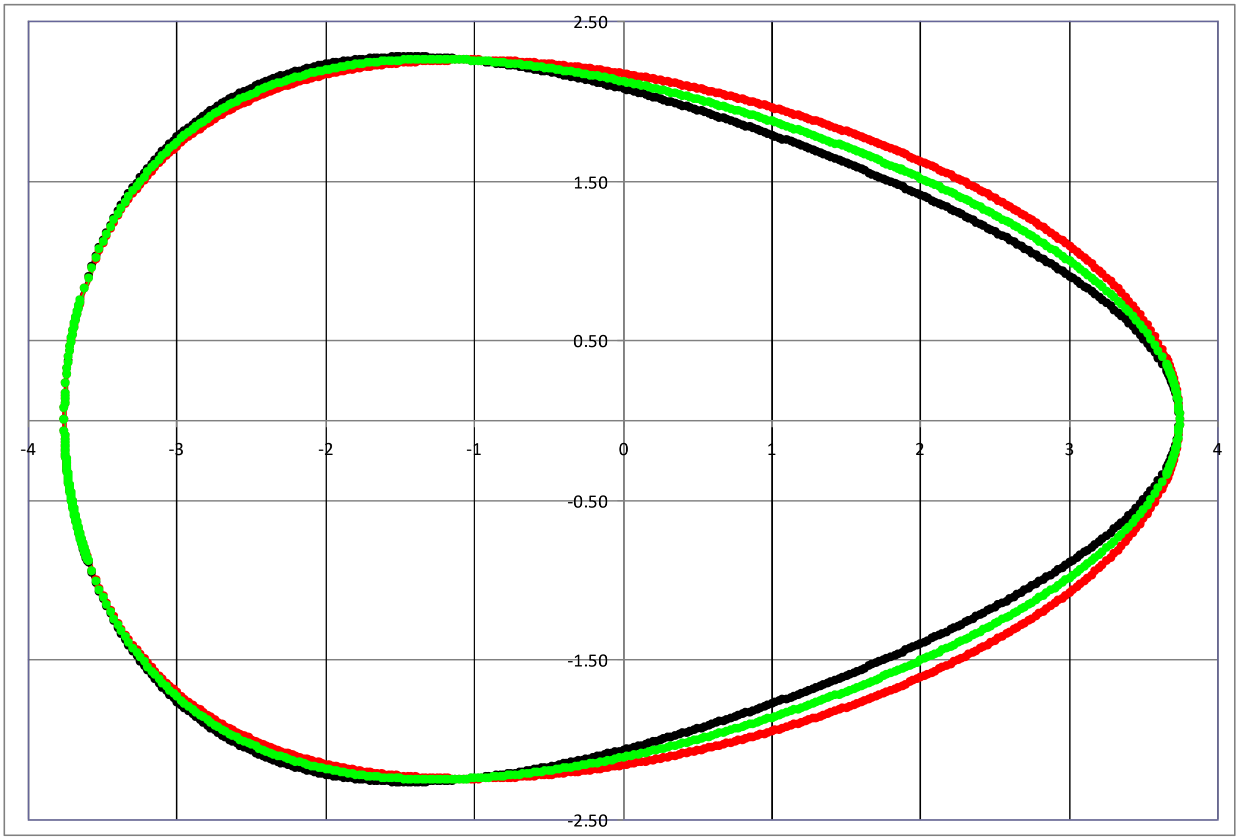


***Figure S3.*** The egg contours according to (EqnS3.9) with $k = 0$ (red); $k = 0.5$ (green) and $k = 1$ (black).

An additional question that required further resolution was how to determine the value of *k* for any tested egg. For this purpose, we considered a characteristic point at $x = L/4$ (***Figure 4***) that can be measured directly. As found in our previous study (***Narushin et al., 2020b***), this point was one of the mostly informative in predicting the parameter *w* in Hügelschäffer's formula. Then, inputting $x = L/4$ into Hügelschäffer's model (Eqn1) and the pyriform formula (EqnS2.21, or Eqn3), we derived the meaning of $y_{L/4}$ at this point respectively for both models:

$$y_{H_{L/4}} = \frac{\sqrt{3}BL}{4\sqrt{L^2 + 2wL + 4w^2}} \qquad \text{(EqnS3.10)}$$

$$y_{c_{L/4}} = \frac{\sqrt{3}BL}{2\sqrt{5.5L^2 + 11Lw + 4w^2}} \qquad \text{(EqnS3.11)}$$

Consequently, the difference between EqnS3.10 and EqnS3.11 leads to the meaning of $\Delta y_{L/4}$ at the point of $x = L/4$:

$$\Delta y_{L/4} = \frac{\sqrt{3}BL}{4} \cdot \frac{\sqrt{5.5L^2 + 11Lw + 4w^2} - 2\sqrt{L^2 + 2wL + 4w^2}}{\sqrt{(L^2 + 2wL + 4w^2)(5.5L^2 + 11Lw + 4w^2)}} \qquad \text{(EqnS3.12)}$$

The value $y_{r_{L/4}}$ of an actual egg can be recalculated from direct measurements of the egg diameter, $D_{L/4}$, at the point of *L*/4 using a caliper as proposed, for example, elsewhere (***Smart, 1991***), machine vision (***Narushin et al., 2020a,b***) or any other proper technique. Taking into consideration that:

$$y_{r_{L/4}} = \frac{D_{L/4}}{2}, \qquad \text{(EqnS3.13)}$$

$$\Delta y_{r_{L/4}} = y_{H_{L/4}} - \frac{D_{L/4}}{2} = \frac{\sqrt{3}BL}{4\sqrt{L^2 + 2wL + 4w^2}} - \frac{D_{L/4}}{2} \qquad \text{(EqnS3.14)}$$

in which $\Delta y_{r_{L/4}}$ is the difference of the *y* values at the point of *x* = *L*/4 of Hügelschäffer's model and an actual egg. Then, the value of the coefficient *k* can be determined using a division of EqnS3.14 and EqnS3.12:

$$k = \frac{\Delta y_{r_{L/4}}}{y_{H_{L/4}}} = \frac{\sqrt{5.5L^2 + 11Lw + 4w^2} \cdot (\sqrt{3}BL - 2D_{L/4}\sqrt{L^2 + 2wL + 4w^2})}{\sqrt{3}BL(\sqrt{5.5L^2 + 11Lw + 4w^2} - 2\sqrt{L^2 + 2wL + 4w^2})} \qquad \text{(EqnS3.15)}$$

Finally, inputting EqnS3.15 into EqnS3.9, we can state that we have obtained the universal model applicable for any avian egg as follows:

$$y = \pm y_H \left(1 - \frac{\sqrt{5.5L^2 + 11Lw + 4w^2} \cdot (\sqrt{3}BL - 2D_{L/4}\sqrt{L^2 + 2wL + 4w^2})}{\sqrt{3}BL(\sqrt{5.5L^2 + 11Lw + 4w^2} - 2\sqrt{L^2 + 2wL + 4w^2})} \cdot \right.$$

$$\left. \cdot \left(1 - \sqrt{\frac{L(L^2 + 8wx + 4w^2)}{2(L - 2w)x^2 + (L^2 + 8Lw - 4w^2)x + 2Lw^2 + L^2w + L^3}}\right)\right) \qquad \text{(EqnS3.16)}$$

or expressing $y_H$ with Eqn1

$$y = \pm \frac{B}{2}\sqrt{\frac{L^2 - 4x^2}{L^2 + 8wx + 4w^2}} \cdot \left(1 - \frac{\sqrt{5.5L^2 + 11Lw + 4w^2} \cdot (\sqrt{3}BL - 2D_{L/4}\sqrt{L^2 + 2wL + 4w^2})}{\sqrt{3}BL(\sqrt{5.5L^2 + 11Lw + 4w^2} - 2\sqrt{L^2 + 2wL + 4w^2})} \cdot \right.$$

$$\left. \cdot \left(1 - \sqrt{\frac{L(L^2 + 8wx + 4w^2)}{2(L - 2w)x^2 + (L^2 + 8Lw - 4w^2)x + 2Lw^2 + L^2w + L^3}}\right)\right) \qquad \text{(EqnS3.17)}$$

**S3 Appendix References**

**Narushin VG**, Lu G, Cugley J, Romanov MN, Griffin DK. 2020a. A 2-D imaging-assisted geometrical transformation method for non-destructive evaluation of the volume and surface area of avian eggs. *Food Control* **112**:107112. doi: 10.1016/j.foodcont.2020.107112.

**Narushin VG**, Romanov MN, Lu G, Cugley J, Griffin DK. 2020b. Digital imaging assisted geometry of chicken eggs using Hügelschäffer's model. *Biosystems Engineering* **197**:45–55. doi: 10.1016/j.biosystemseng.2020.06.008.

**Smart IHM**. 1991. Egg-shape in birds. In: Deeming DC, Ferguson MWJ, editors. Egg Incubation: Its Effects on Embryonic Development in Birds and Reptiles. Cambridge: Cambridge University Press. pp. 101–116. doi: 10.1017/CBO9780511585739.009.

407
